# Use of vivo-morpholinos for gene knockdown in the postnatal shark retina

**DOI:** 10.1101/2022.07.11.499558

**Authors:** Mariña Rodríguez-Arrizabalaga, Ismael Hernández-Núñez, Eva Candal, Antón Barreiro-Iglesias

## Abstract

Work in the catshark *Scyliorhinus canicula* has shown that the evolutionary origin of postnatal neurogenesis in vertebrates is earlier than previously thought. Thus, the catshark can serve as a model of interest to understand postnatal neurogenic processes and their evolution in vertebrates. One of the best characterized neurogenic niches of the catshark CNS is found in the peripheral region of the retina. Unfortunately, the lack of genetic tools in sharks limits the possibilities to deepen in the study of genes involved in the neurogenic process. Here, we report a method for gene knockdown in the juvenile catshark retina based on the use of Vivo-Morpholinos. To establish the method, we designed Vivo-Morpholinos against the proliferation marker PCNA. We first evaluated the possible toxicity of 3 different intraocular administration regimes. After this optimization step, we show that a single intraocular injection of the PCNA Vivo-Morpholino decreases the expression of PCNA in the peripheral retina, which leads to reduced mitotic activity in this region. This method will help in deciphering the role of other genes potentially involved in postnatal neurogenesis in this model.

**Summary statement:** In this study, we report the development of a method for the use of Vivo-Morpholinos in the postnatal shark retina, which will allow to decipher the role of different genes in retinal neurogenesis in sharks.

## 1 INTRODUCTION

Neurogenesis is the process by which progenitor cells generate new neurons. During development and ageing, this process is progressively restricted to the so-called neurogenic niches, where stem cell self-renewal occurs. Postnatal neurogenic niches and cell proliferation/neurogenesis in the central nervous system (CNS) are more widespread and abundant in fish than in mammals, which facilitates the study of postnatal/adult neurogenesis in this vertebrate group. Although most of the knowledge of postnatal neurogenesis in fish comes from teleost fish, recent studies in chondrichthyan fish suggest that the evolutionary origin of this process is earlier than previously thought (Docampo-Seara et al., 2020). Chondrichthyes are the oldest extant gnathostome vertebrates and their key phylogenetic position allows to find characters that were fixed prior to the gnathostome radiation. Indeed, recent work of our group using the lesser spotted dogfish or catshark (*Scyliorhinus canicula*) as a model is providing interesting comparative information about the postnatal neurogenic process in the CNS of vertebrates (Docampo-Seara et al., 2020; Hernández-Núñez et al., 2021b). For example, in catshark telencephalic neurogenic niches, different subtypes of progenitor cells like radial glial progenitor cells, intermediate progenitor-like cells and migrating neuroblasts have been described based on the expression of typical (and evolutionary conserved) markers of each of these progenitor cell types (Docampo-Seara et al., 2020).

Some of the best characterized neurogenic niches in the CNS of fish are also found in the retina (reviewed in Amato et al., 2004; Moshiri et al., 2004; Ail and Perron, 2017; Miles and Tropepe, 2021) and, again, work in teleost models has provided important information on the genes and molecular pathways controlling postnatal neurogenesis from progenitor cells in these niches (e. g., Conner et al., 2014). In catsharks, as in teleost and other fish usually used as models to study retinal neurogenesis, the peripheral postnatal retina contains a circumferential ring of proliferating cells located between the ciliary epithelium and the mature central retina known as ciliary marginal zone (CMZ) (Sánchez-Farías and Candal 2015, 2016; Ferreiro-Galve et al., 2021; Hernández-Núñez et al., 2021b). The large size of the retina and the slow pace of retinal development in catshark as compared to that of teleost fish allowed us to identify a transition zone (TZ) located between the CMZ and the central retina. This TZ, which had been overlooked in other fish species, contains different types of progenitor cells, like neuroepithelial-like cells, different types of radial glia-like cells and migrating neuroblasts (Ferreiro-Galve et al., 2010a; Sánchez-Farías and Candal, 2015, 2016; Hernández-Núñez et al., 2021b). Recent transcriptomic retinal data from *S. canicula* revealed several genes whose expression changes between juveniles and adults and that could be involved in maintaining a high proliferative and neurogenic activity in the juvenile retina of this species (Hernández-Núñez et al., 2021b). These data provide an excellent resource to identify new genes and signaling pathways controlling neurogenesis in the vertebrate retina. This, together with its advantages over other fish models (see above), make the catshark retina an important model for future functional work in this field. Unfortunately, the lack of stable transgenic or mutant shark lines limits the possibilities to deepen in the study of genes involved in the neurogenic process of the catshark retina.

Morpholino oligonucleotides (MOs) are one of the most widely used anti-sense knockdown techniques for blocking gene expression. MOs are chemically synthetised oligomers that are typically constituted by 25 bases that are base-pairing complementary to the target RNA. MOs have been used to inhibit the translation of RNA transcripts i*n vivo* by modification of mRNA splicing, mRNA stability and translation (reviewed by Bill et al., 2009). The specificity in the recognition between the MO and its complementary mRNA sequence allows a high affinity and a low level of secondary effects. For example, microinjection of MOs has been used for many years in loss of function experiments in developing zebrafish (reviewed in Wang and Cao, 2021). However, MOs show low cell penetration ability, which limits their use to early developmental stages [1- to 8-cell-stage embryos (reviewed in Bill et al., 2009)]. But more recently, a new MO variant has been developed, the so-called Vivo-MOs. Vivo-MOs are MOs linked to a guanidinium dendrimer, which allows penetration of the Vivo-MO in cells from cell culture medium, blood, or cerebrospinal fluid (Morcos et al., 2008). For example, Vivo-MOs have already been used for gene knockdown in the mouse retina via intra-vitreal injections (Owen et al., 2012). Thus, Vivo-MOs open big possibilities for the manipulation of gene expression in postnatal individuals and especially in non-conventional animal models (like sharks) in which genetically modified specimens are not yet available.

Here, our aim was to establish a method for the use of Vivo-MOs to knockdown gene expression in the retina of *S. canicula* juveniles. Since the juvenile retina shows high proliferative and mitotic activity in the CMZ, we decided to use a translation blocking Vivo-MO generated against the proliferating cell nuclear antigen (PCNA). PCNA is a proliferation marker that is expressed during the cell cycle (G1, S and G2 phases; Zerjatke et al., 2017) and that shows very high expression in the peripheral retina of juvenile catsharks (Ferreiro-Galve et al., 2010a; Hernández-Núñez et al., 2021b). Our data show that a single intraocular injection of a PCNA Vivo-MO significantly decreases PCNA expression in the CMZ and that this leads to a decrease in mitotic activity (as shown by pH3 immunolabelling). To our knowledge, this is the first study to report a gene manipulation method in sharks. Designing a method for the use of Vivo-MOs in the catshark retina will help in deciphering the role of other genes potentially involved in postnatal neurogenesis in this species.

## 2 RESULTS AND DISCUSSION

### 2.1 Assessment of Vivo-MO toxicity

In this study, we generated a PCNA translation blocking Vivo-MO targeting the 5’untranslated region of the catshark PCNA mRNA (Fig. 1A) and used the standard control Vivo-MO from Gene Tools as a control. A Blastn (Stephen et al., 1997) search of the PCNA Vivo-MO sequence in the *S. canicula* genome (sScyCan1.1 reference, Annotation Release 100, GCF_902713615.1) showed that only a maximum of 14 bases of the Vivo-MO can be aligned with other gene sequences, which suggest high specificity (not shown). For example, 5-base mismatch MO controls are sometimes used as negative controls in MO experiments (Stainier et al., 2017) and in this case the minimum mismatch with other genes would be of 11 bases.

**FIGURE 1.**
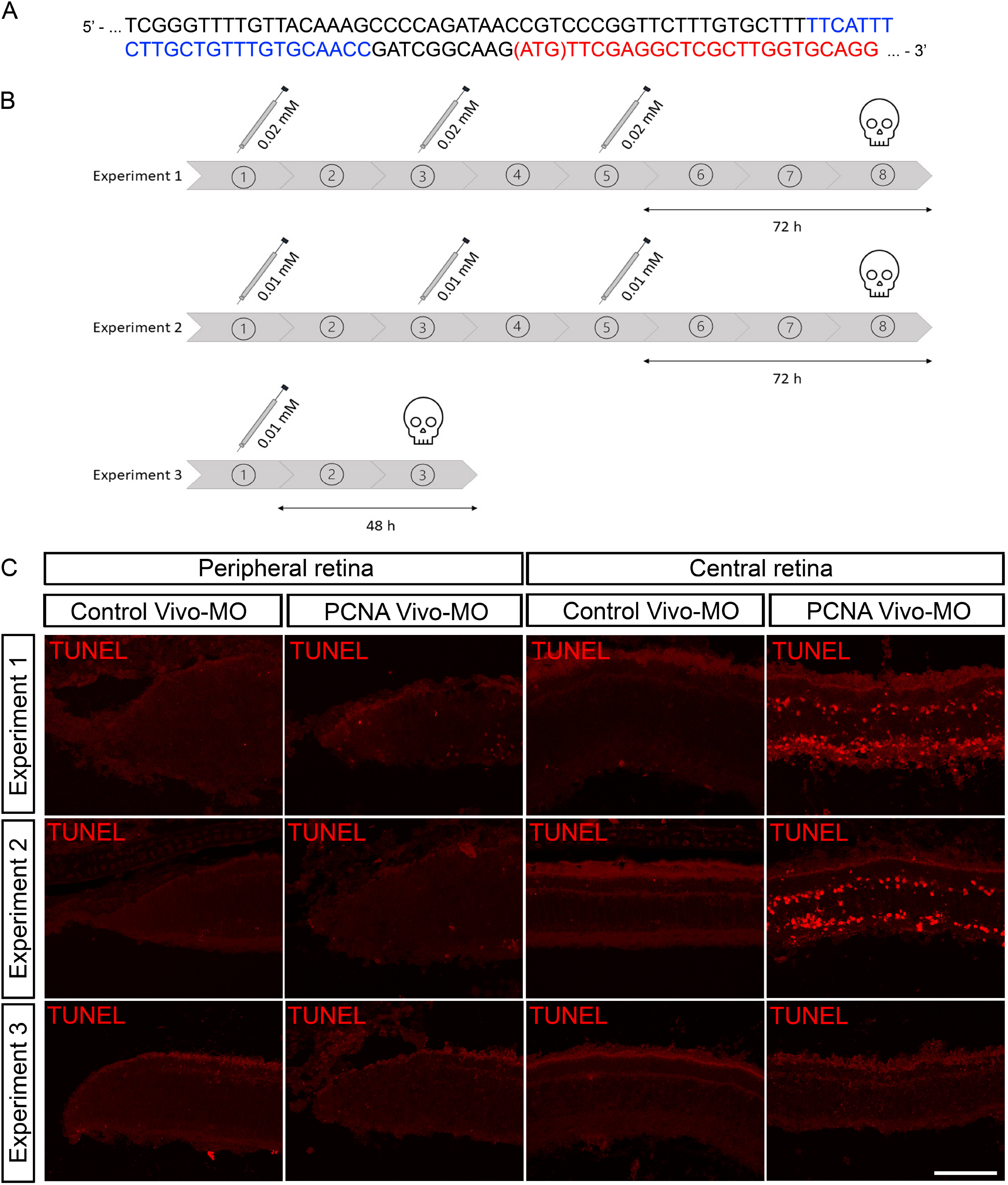
(A) Partial sequence of the catshark PCNA mRNA. Protein coding sequence in shown in red and 5’ untranslated sequence in black. The target sequence of the PCNA Vivo-MO in the 5’ untranslated sequence is indicated in blue. The ATG codon is indicated between parentheses. (B) Diagram of the experimental design for experiments 1 to 3. (C) Photomicrographs of TUNEL labelling in peripheral and central retinas after administration of the Control Vivo-MO or the PCNA Vivo-MO in each experiment. Scale bar: 100 μm.

Vivo-MOs are considered as a useful, specific, and efficient anti-sense knockdown tool with usually little to no toxicity after treatment; however, problems with toxicity have also been described in some studies (reviewed by Ferguson et al., 2014). Because of this, we first decided to test 3 different regimes of intraocular Vivo-MO administration with different number of injections, concentrations of Vivo-MOs and time points of analysis (Experiment 1: 3 doses of 15 μL at a concentration 0.02 mM, n = 5; Experiment 2: 3 doses of 15 μL at a concentration of 0.01 mM, n = 2; Experiment 3: 1 dose at a concentration 0.01 mM, n = 11; Fig. 1B). To test the potential toxicity of the Vivo-MOs, we used Tdt-mediated dUTP Nick End Labelling (TUNEL) to detect the presence of apoptotic nuclei. The Control Vivo-MO was injected into the right eye and the PCNA Vivo-MO into the left eye of each specimen. The presence of a few TUNEL positive cells has been previously detected in postnatal retinas of *S. canicula* (Bejarano-Escobar et al., 2013). An increase in TUNEL labelling above background levels was never detected in the peripheral or central retinas that received the Control Vivo-MO in any of the 3 experimental conditions (Fig. 1C). Cell death was not increased in the peripheral retina after PCNA Vivo-MO administration in any of the experimental conditions (Fig. 1C). In contrast, we observed a clear increase in apoptotic cell death in the central retina of eyes that received 3 injections of the PCNA Vivo-MO (experiments 1 and 2; Fig. 1C). No increase in cell death was detected in the peripheral and central retinas after a single intraocular injection of the PCNA Vivo-MO (experiment 3; Fig. 1C).

The presence of apoptotic cells in the central retina in animals that received 3 injections of the PCNA Vivo-MO (experiments 1 and 2) could be explained by a cumulative toxic effect of the PCNA Vivo-MO in cells of the central retina that normally contains very few cells showing PCNA expression (Hernández-Núñez et al., 2021b). We did not observe an increase in cell death in the peripheral retina [which shows high numbers of PCNA+ cells (Hernández-Núñez et al., 2021b; see below)] with the PCNA Vivo-MO, even in experiments 1 and 2, which suggests that apoptotic cell death in the central retina could be caused by the simple accumulation of unbound PCNA Vivo-MO. However, the fact that the administration of the Control Vivo-MO did not lead to increased cell death suggests that toxicity with the PCNA Vivo-MO could be related to higher chances of off-target effects (see Eisen and Smith, 2008). For example, in zebrafish embryos between 15 and 20% of the MOs can activate p53-induced apoptotic cell death in neurons (Robu et al., 2007; see Eisen and Smith, 2008). In any case, our TUNEL labelling data shows that a single dose of the PCNA Vivo-MO (15 μL at a concentration of 0.01 mM) did not cause an increase in cell death in the retina and, therefore, this administration regime was selected for subsequent analyses (see below).

### 2.2 Effect of the PCNA Vivo-MO on PCNA expression and cell proliferation

After optimising the Vivo-MO administration regime we decided to analyse whether the single intraocular administration of the PCNA-Vivo MO was able to knockdown the expression of PCNA in the peripheral retina (CMZ+TZ). As previously reported during normal development and in early postnatal life (Ferreiro-Galve et al., 2010a; Sánchez-Farías and Candal, 2015, 2016; Hernández-Núñez et al., 2021b), in the retina of juveniles, most PCNA expression was found throughout the CMZ and TZ (Fig. 2A, B). To quantify changes in PCNA expression we measured both the area showing PCNA immunoreactivity in the peripheral retina (Fig. 2C) and the mean fluorescence intensity of PCNA immunoreactivity in the peripheral retina (Fig. 2D). Quantification of the area showing PCNA labelling revealed no significant differences between the Control Vivo-MO retinas and those that received the PCNA Vivo-MO (Fig. 2A-C). However, quantifications of mean PCNA fluorescence intensity revealed a significant decrease in fluorescence intensity (PCNA expression) after PCNA Vivo-MO administration (Fig. 2A, B, D). These results indicate that the administration of the PCNA Vivo-MO can knockdown the expression of PCNA, but it does not completely block its expression (which would have been detected as a reduction in the area occupied by PCNA+ cells).

**FIGURE 2.**
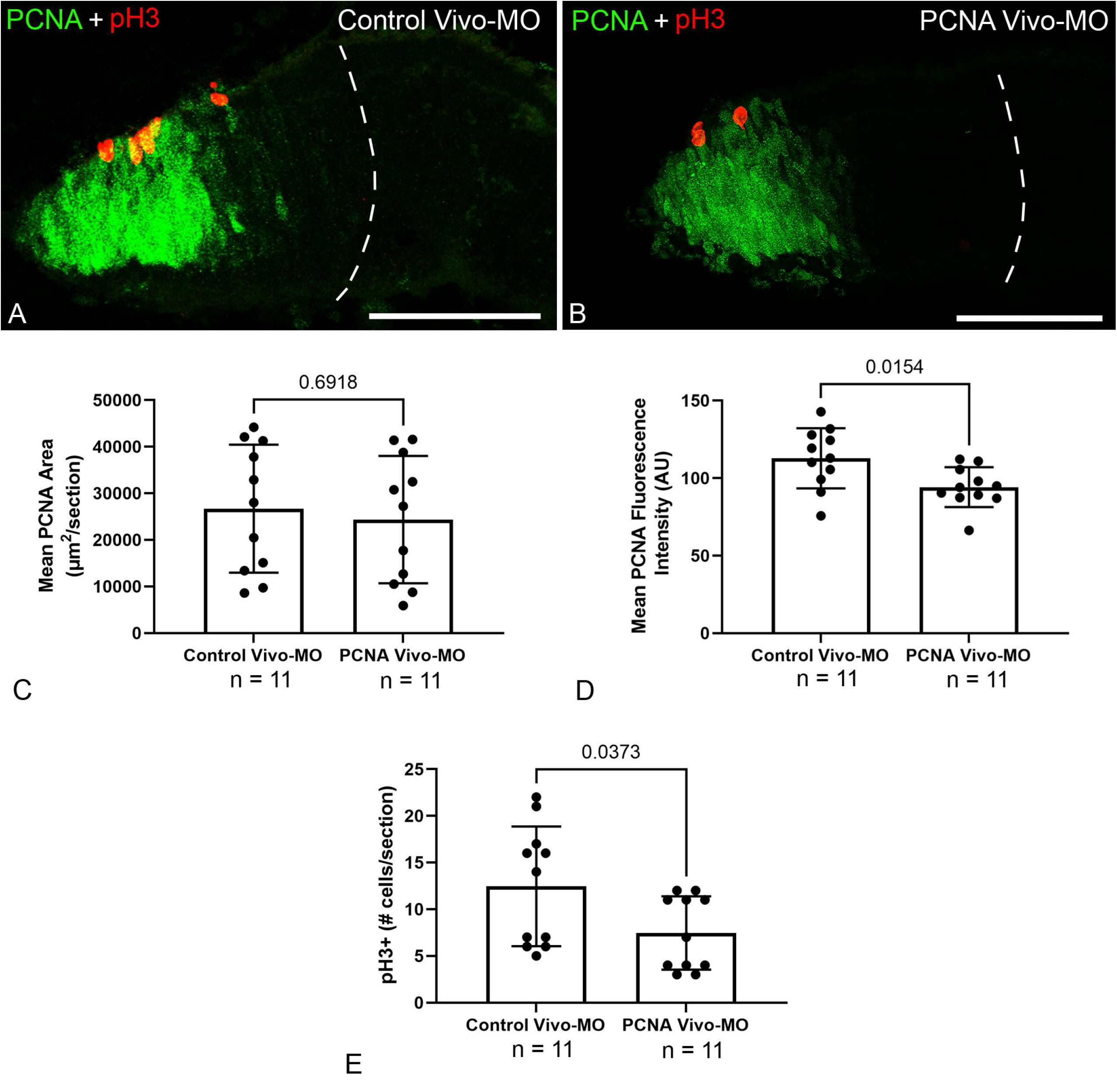
The PCNA Vivo-MO decreases PCNA expression and the number of mitotic cells (pH3+) in the peripheral retina. (A) Photomicrograph of the peripheral retina after Control Vivo-MO administration. (B) Photomicrograph of the peripheral retina after PCNA Vivo-MO administration. The dashed lines indicate the limit between the peripheral and the central retinas. (C) Graph showing the lack of significant change (unpaired t-test) in the PCNA+ area between Control Vivo-MO retinas (26698 ± 4140 μm^2^) and PCNA Vivo-MO retinas (24350 ± 4118 μm^2^). (D) Graph showing a significant decrease (unpaired t-test) in mean fluorescence intensity (arbitrary units) in the peripheral retina between Control Vivo-MO retinas (112.8 ± 5.853) and PCNA Vivo-MO retinas (94.21 ± 3.884). (E) Graph showing a significant decrease (Mann-Whitney U test) in the number of pH3+ cells in the peripheral retina between Control Vivo-MO retinas (12.45 ± 1.932 cells per section) and PCNA Vivo-MO retinas (7.455 ± 1.186 cells per section). Scale bars: 100 μm.

The lack of a complete blockage of PCNA translation could be a limitation to the use of Vivo-MO PCNA for loss-of-function studies. To explore to what extent decreasing PCNA levels in each cell is sufficient to affect cell division, we decided to quantify the number of mitotic (pH3+) cells in this region. pH3+ cells were mostly found restricted to the ventricular (apical) region of the peripheral retina (Fig. 2A, B). Cell quantifications revealed a statistically significant decrease in the number of pH3+ cells after PCNA-Vivo-MO administration (Fig. 2A, B, E). Overall, our data shows a successful knockdown of PCNA expression in the peripheral retina of catshark juveniles after a single intraocular injection of the PCNA Vivo-MO and that the decrease in PCNA expression leads to reduced mitotic activity in this region. These results validate our methodological design for the future use of Vivo-MOs to knockdown gene expression in the catshark retina and modify the behaviour of progenitor cells, though the dosage and schedule of administration of different Vivo-MOs might need to be adjusted on a case-by-case basis. MOs had been previously applied to the adult zebrafish retina, but in a process that involved electroporation after the MO injection into the vitreous (Thummel et al., 2008, 2011) or cutting the optic nerve for delivery to retinal ganglion cells (Ogai et al., 2014). The use of Vivo-MOs will facilitate the use and application of this gene knockdown technique to fish models.

Interestingly, when looking at our quantitative data in Control Vivo-MO retinas we noticed that there could be a possible negative correlation between body size and the amount of proliferative activity in the peripheral retina (see Table 1 with the specimens ordered by body length). So, in subsequent analyses we decided to analyse this correlation and possible differential effects of the PCNA Vivo-MO in younger/shorter and older/longer juvenile catsharks (with high and low cell proliferation levels, respectively).

**Table 1.**
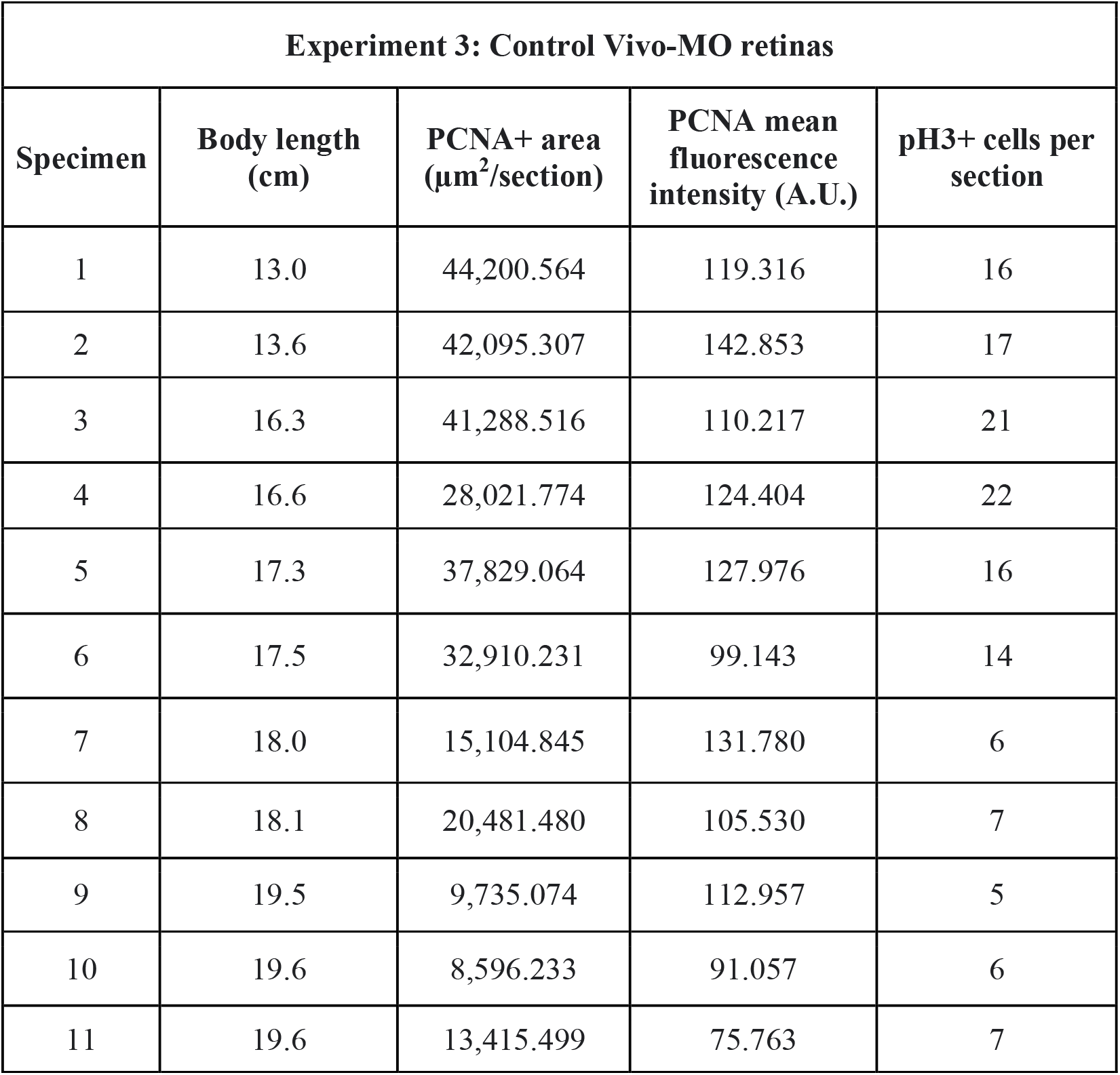
Body length of specimens in experiment 3 with proliferation values for each Control Vivo-MO retina. Note that the specimens were ordered based on their body length.

### 2.3 Proliferative/mitotic activity and body length

First, we carried out statistical correlation analyses to confirm the possible correlation between an increase in body length/age and a decrease in cell proliferation in the peripheral retina (Fig. 3) of the eyes that received the Control Vivo-MO. Simple linear regression analyses revealed a significant correlation between body length for each specimen and values of PCNA area (Fig. 3A-C), PCNA fluorescence intensity (Fig. 3A, B, D), and number of pH3+ cells (Fig. 3A, B, E). These significant correlations show that with increased age/body length there is a decrease in proliferative/mitotic activity in the retina. This coincides well with previous data from our group showing a loss of proliferative activity between catshark juveniles and sexually mature adults (Hernández-Núñez et al., 2021b) and indicates that the loss of proliferative activity starts before sexual maturation during the juvenile stage. A decline in proliferative activity during ageing has been also observed in the zebrafish retina (Van Houcke et al., 2019; Hernández-Núñez et al., 2021a).

**FIGURE 3.**
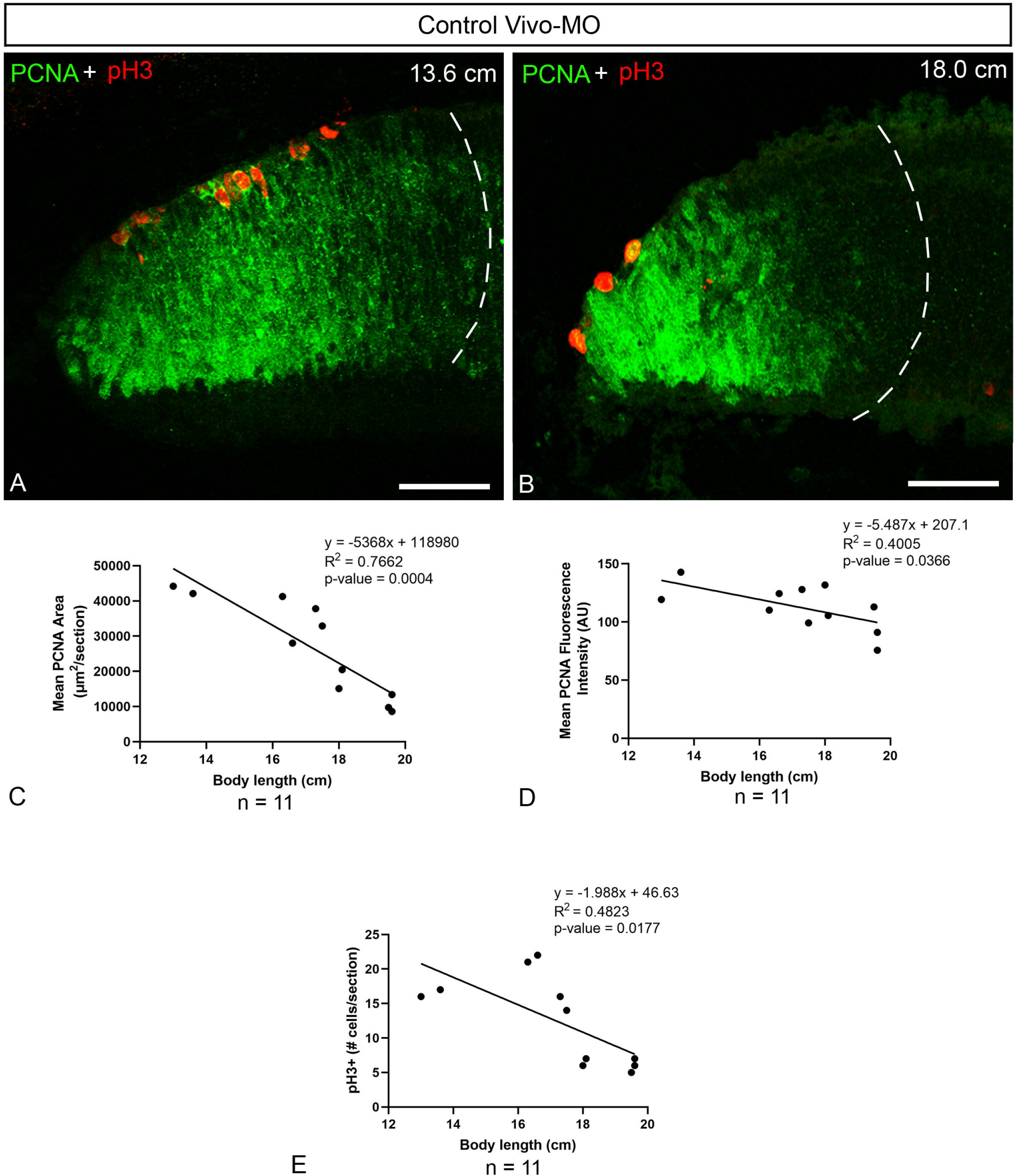
Negative correlation between PCNA+ area, PCNA fluorescence intensity or number of pH3+ cells of Control Vivo-MO retinas and body length. (A) Photomicrograph showing the expression pattern of PCNA and pH3 in the peripheral retina of a specimen of 13.6 cm. (B) Photomicrograph showing the expression pattern of PCNA and pH3 in the peripheral retina of a specimen of 18.0 cm. Dashed lines indicate the limit between the peripheral and central retina. (C) Graph showing simple linear regression of fluorescence intensity with respect to body length. (D) Graph showing simple linear regression of the PCNA+ area with respect to body length. (E) Graph showing simple linear regression of the number of pH3+ cells with respect to body length. For individual values see Table 1. Scale bars: 50 μm.

When looking at the correlation graphs there is a clear decrease in mitotic activity (pH3) in individuals with a body length of more than 17 cm (Fig. 3E). Based on this, we decided to reanalyse the effect of the PCNA Vivo-MO separately for the specimens between 13 and 16.6 cm in body length and specimens with a body length between 17.3 and 19.3 cm.

As in the analyses that included all the specimens (see section 2.2), we did not detect significant changes in the area occupied by PCNA immunoreactivity in the 13-16.3 cm (not shown; Control Vivo-MO: 38902 ± 3678 μm^2^; PCNA Vivo-MO: 32312 ± 2410 μm^2^; p = 0.1846, unpaired t test) or in the 17.3-19.3 cm (not shown; Control Vivo-MO: 19725 ± 4329 μm^2^; PCNA Vivo-MO: 19799 ± 5767 μm^2^; p = 0.9015, Mann-Whitney U test) groups after PCNA Vivo-MO administration. However, PCNA fluorescence intensity and numbers of pH3+ cells were significantly decreased after the administration of the PCNA Vivo-MO in the 13-16.3 cm group (Fig. 4A, B, E, G) and not in juveniles of the 17.3-19.3 cm group (Fig. 4C, D, F, H). The reduction in mitotic activity (pH3 labelling) was even more significant in the 13-16.3 cm group (n = 4; Fig. 4G) than with all the specimens (n = 11; Fig. 2E; see above). These results indicate that this administration regime is more effective in younger/shorter juvenile catsharks and that the manipulation of postnatal proliferative and neurogenic processes will be facilitated in young postnatal juveniles because of their higher rates of cell proliferation.

**FIGURE 4.**
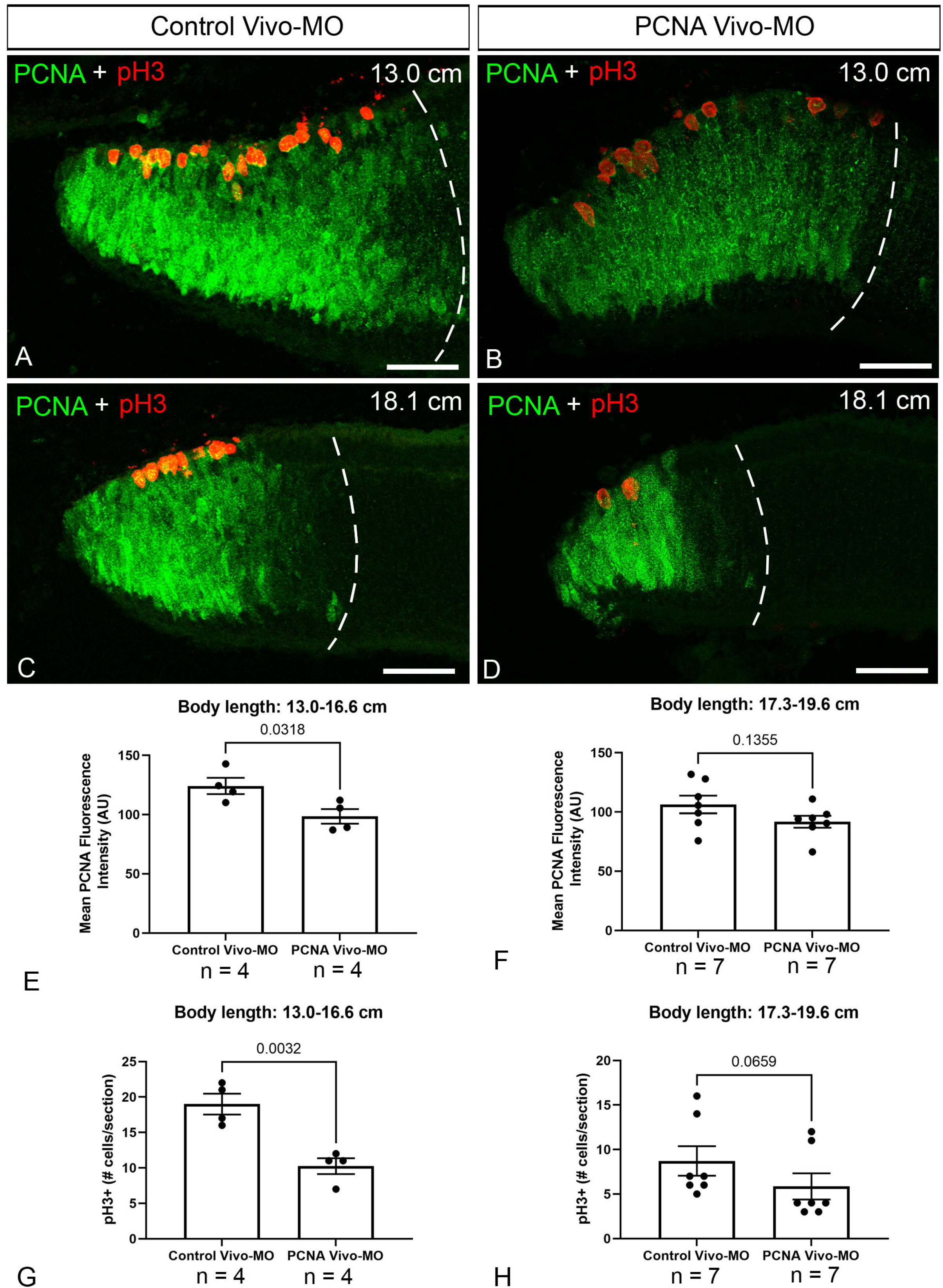
Changes in mean PCNA fluorescence intensity and in the number of mitotic pH3+ cells in Control Vivo-MO and PCNA Vivo-MO retinas coming from juvenile specimens of short (13.0-16.6 cm) or long (17.3-19.6) body length. (A-D) Photomicrographs of Control Vivo-MO or PCNA Vivo-MO peripheral retinas in specimens of the 13.0-16.6 cm group or the 17.3-19.6 cm group. Dashed lines indicate the limit between the peripheral and central retinas. (E) Graph showing a significant decrease (unpaired t-test) in mean PCNA fluorescence intensity (A.U.) between Control Vivo-MO (124.2 ± 6.676) and PCNA Vivo-MO (98.51 ± 6.154) peripheral retinas in specimens of the 13.0-16.6 cm group. (F) Graph showing the lack of significant changes (unpaired t-test) in mean PCNA fluorescence intensity (A.U.) between Control Vivo-MO (106.3 ± 7.537) and PCNA Vivo-MO (91.75 ± 5.091) peripheral retinas in specimens of the 17.3-19.6 cm group. (G) Graph showing a significant decrease (unpaired t-test) in the number of pH3+ cells between Control Vivo-MO (19.00 ± 1.472 cells per section) and PCNA Vivo-MO (10.25 ± 1.109 cells per section) peripheral retinas in specimens of the 13.0-16.6 cm group. (H) Graph showing the lack of significant changes (Mann-Whitney U test) in the number of pH3+ cells between Control Vivo-MO (8.714 ± 1.658 cell per section) and PCNA Vivo-MO (5.857 ± 1.471 cells per section) peripheral retinas in specimens of the 17.3-19.6 cm group. Scale bars: 100 μm.

In conclusion, our work provides a methodological basis for future manipulation of genes of interest in the analysis of postnatal neurogenic processes in the catshark retina. For example, recent transcriptomic data from our group revealed that signalling pathways like the Notch, Shh, Robo/Slit or Wnt pathways could play a role in regulating cell proliferation and neurogenesis in the postnatal *S. canicula* retina (Hernández-Núñez et al., 2021b). Vivo-MOs can complement the use of pharmacological tools to manipulate these pathways or even allow the manipulation of non-druggable genes.

We must be also aware of the limitations of this technique and future work should also try to improve the available tools/controls for Vivo-MO use in sharks. For example, guidelines proposed for morpholino use in zebrafish suggest a series of extra controls (apart from the use of control MOs or the use of antibodies to detect protein knockdown as shown here) when performing MO experiments, which could include: 1) a comparison of the morphant phenotype to that of a mutant; 2) mRNA rescue by administration of an mRNA lacking the MO-binding site; or 3) MO injection in homozygous mutants for the target gene (Stainier et al., 2017). These methods are not yet available in sharks (especially in the case of mutant lines), but a first step could be to try to find a suitable way (e.g., viral vectors or electroporation) to deliver mRNAs or RNAs for CRISPR interference to the catshark retina. Neuroscience research focuses largely on a handful of animal models, but reaching beyond the usual models (rodents, worms, flies or zebrafish) will allow us to identify common principles in neural development and function. Expanding the methodological tools will allow to work on an expanded range of animal models for the study of neurogenesis.

## 3 MATERIAL AND METHODS

### 3.1 Animals

Juveniles (n = 18; 11.2 to 19.6 cm long) of both sexes of *S. canicula* were kindly provided by the aquarium Finisterrae (A Coruña, Spain) and kept in seawater tanks under standard conditions of temperature (15-16 °C), pH (7.5-8.5), and salinity (35 g/L). All experimental procedures were performed following the guidelines established by the European Union (2010/63/EU) and the Spanish Royal Decree 1386/2018 for the care and handling of animals in research and were approved by the Bioethics Committee of the University of Santiago de Compostela (license number 15004/2022/001).

### 3.2 Vivo-MO intraocular administration

Dilutions of Vivo-MOs (0.02 mM or 0.01 mM) were obtained by resuspending the lyophilized stocks in Milli-Q water. Animals were anesthetised with 0.01 g tricaine methanesulfonate (MS-222, Sigma, St. Louis, MO) in 200 mL of seawater. Then, 15 μL of the solutions containing the PCNA Vivo-MO [sequence: 5’-GGTTGCACAAACAGCAAGAAATGAA - 3’; designed by Gene Tools LLC (Philomath, OR, United States) based on the *S. canicula* mRNA PCNA sequence (GenBank reference XM_038785554.1)] were injected into the left eye. 15 μL of the solutions containing the Control Vivo-MO (Gene Tools Standard Control Vivo-MO; sequence: 5’-CCTCTTACCTCAGTTACAATTTATA-3’) were injected into the right eye. Injections in both eyes were performed through the cornea and in the intravitreal space with a sterile syringe and a 30G needle. After the injections, animals were left to recover in individual tanks in 200 mL of aerated seawater. The number of injections and timing of perfusion/fixation for each experimental condition (experiments 1 to 3) can be found in Figure 1A.

### 3.3 Tissue Preparation and Histology

At the end of the experiments, animals were deeply anaesthetised with 0.1 g MS-222 in 200 mL of seawater and then perfused intracardially with elasmobranch Ringer’s solution (1.7 % NaCl, 0.024 % KCl, 0.031 % CaCl_2_, 0.044 % MgCl_2_, 0.113 % Na_2_SO_4_, 0.049 % NaCO_3_H, 2.7 % urea; see Ferreiro-Galve et al., 2008) followed by perfusion with 4 % paraformaldehyde (PFA) in 0.1 M phosphate buffer (PB) containing 1.75 % urea (elasmobranch PB; pH 7.4). The eyes were removed and postfixed in 4 % PFA for 2 days at 4 °C. After rinsing in phosphate buffered saline (PBS), the eyes were cryoprotected with successive solutions of 10 %, 20 % and 30 % sucrose in PBS, embedded in Neg-50TM (Thermo Scientific, Kalamazoo, MI), and frozen with liquid nitrogen-cooled isopentane. Four parallel series of transvers sections (18 μm thick) were obtained on a cryostat and mounted on Superfrost Plus slides (Menzel-Glässer®, Madison, WI, USA).

### 3.4 Immunofluorescence

Sections were first pre-treated with 0.01 M citrate buffer pH 6.0 for 30 min at 90 °C for heat-induced epitope retrieval, allowed to cool for 20 min at room temperature and rinsed in tris buffered saline (TBS; pH 7.4) for 5 min. Then, sections were incubated overnight at room temperature with a combination of 2 different primary antibodies: a mouse monoclonal anti-PCNA antibody (1:500; Sigma-Aldrich; catalogue number P8825; RRID: AB_477413) and a rabbit polyclonal anti-pH3 antibody (1:300; Millipore; Billerica; MA, USA; catalogue number 06-570; RRID: AB_310177). Sections were rinsed 3 times in TBS for 10 min each and incubated for 1 h at room temperature with a combination of 2 fluorescent dye-labelled secondary antibodies: a Cy3-conjugated goat anti-rabbit antibody (1:200; Invitrogen, Waltham, MA, USA; catalogue number A10520) and a FITC-conjugated goat anti-mouse antibody (1:200; Invitrogen; catalogue number F2761). All antibody dilutions were made in TBS containing 15 % normal goat serum (Millipore), 0.2 % Triton X-100 (Sigma-Aldrich), and 2 % bovine serum albumin (Sigma-Aldrich). Then sections were rinsed 3 times in TBS for 10 min each and in distilled water for 30 min, allowed to dry for 30 min at 37 °C, and mounted with 100 μL of MOWIOL® 4-88 (Calbiochem, Daemstadt, Germany).

### 3.5 Specificity of Antibodies

PCNA is present in proliferating cells and although its expression is stronger during the S phase, it persists along the entire cell cycle excepting the mitotic period (Zerjatke et al., 2017). The anti-PCNA antibody has been previously used to label progenitor cells in the brain and retina of *S. canicula* (i.e, Quintana-Urzainqui et al., 2014; Ferreiro-Galve et al., 2010a, b, 2012; Sánchez-Farías and Candal, 2015; Hernández-Núñez et al., 2021b). The anti-pH3 antibody has been also widely used in the brain and retina of *S. canicula* as a marker of mitotic cells (Ferreiro-Galve et al., 2010a, b; Bejarano-Escobar et al., 2012; Quintana-Urzainqui et al., 2014; Hernández-Núñez et al., 2021b).

### 3.6 TUNEL labelling

We used the Tdt-mediated dUTP Nick End Labelling (TUNEL) Kit (Roche, Mannheim, Germany; catalogue number 12156792910) to detect apoptotic nuclei. Sections were pre-treated, first with MetOH at −20 °C for 15 min and then with 0.01 M citrate buffer pH 6.0 for 30 min at 90 °C. Then, sections were rinsed 3 times in PBS for 10 min each and incubated with a mixture of 5 μL of enzyme solution (terminal deoxynucleotidyl transferase) and 45 μL of labelling solution (TMR red labelled nucleotides) per slide for 90 min at 37 °C. Then, sections were rinsed 3 times in PBS for 15 min each and 2 times in distilled water for 10 min each, allowed to dry for 30 min at 37 °C, and mounted with 100 μL of MOWIOL® 4-88.

### 3.7 Image Acquisition

Images of fluorescent labelled sections were taken with a Leica Stellaris 8 confocal microscope (Leica Microsystems, Mannheim, Germany) with a combination of blue and green excitation lasers and using a 20x or 40x objectives. Confocal optical sections were taken at steps of 1 μm along the z-axis. Collapsed images of the complete retinal 18 μm sections were generated with the LAS X Software (Leica, Wetzlar, Germany). Confocal images were always taken with the same confocal microscope and acquisition software parameters for retinas coming from Control and PCNA Vivo-MO eyes. For figure preparation, contrast and brightness of the images were minimally adjusted (and always after quantifications) using Adobe Photoshop 2021 (Adobe, San Jose, CA, USA).

### 3.8 Quantifications

We quantified the area showing PCNA+ labelling and measured the mean fluorescence intensity of PCNA+ labelling of the peripheral retina in confocal images (20x objective). The number of mitotic cells (pH3+) in the peripheral retina was quantified under an Olympus fluorescence microscope. The limit between the peripheral retina and the central retina was established based on morphological differences (for example the laminated structure of the central retina) and based on the expression pattern of PCNA (which is mainly found in the peripheral region of the retina).

The area with PCNA+ labelling was quantified using the Measure tool in the Fiji software (Schindelin et al., 2012), in 1 out of each 4 consecutive sections for each retina (only 1 peripheral retina was quantified in each section). Then, we calculated the mean value of PCNA+ area per section for each retina and used that value for statistical analyses (each dot in the graphs represents 1 retina from 1 animal).

The mean fluorescence intensity of PCNA+ labelling was quantified in confocal photomicrographs using the Histogram tool in the Fiji software (Schindelin et al., 2012), in 1 out of each 4 consecutive sections for each retina (only 1 peripheral retina was quantified in each section). Then, we calculated the mean value of PCNA fluorescence intensity per section for each retina and used that value for statistical analyses (each dot in the graphs represents 1 retina from 1 animal).

The number of pH3+ cells were manually counted under the fluorescence microscope in 1 out of each 4 consecutive sections for each retina (only 1 peripheral retina was quantified in each section). Then, we calculated the mean value of pH3+ cells per section for each retina and used that value for statistical analysis (each dot in the graphs represents 1 retina from 1 animal).

### 3.9 Statistical analyses

Statistical analyses were performed with Prism 9 (GraphPad software, La Jolla, CA, USA). Normality of the data in groups with n = 11 was determined with the D’Agostino & Pearson test. For groups with a lower n number (n = 4 or 7), we used the Shapiro-Wilk normality test. To determine statistically significant differences (p ≤ 0.05) between two groups of normally distributed data we used an unpaired (Student’s) t-test (two-tailed). To determine statistically significant differences of non-normally distributed data we used a Mann Whitney U test (two-tailed). To determine the possible correlation between PCNA+ area, mean PCNA fluorescence intensity or number of pH3+ cells with body length we used a simple linear regression. We determined the significance of the slope with respect to zero (p ≤ 0.05) and calculated the equation of the straight line and the R^2^.

## Acknowledgements

We would like to thank the *Servicio de Microscopía* of the University of Santiago de Compostela and Dr. Mercedes Rivas Cascallar for confocal microscope facilities and technical help.

## Data availability

Histological material and raw quantitative data are available from the authors.

